# Field cancerization impacts tumor development, T-cell exhaustion and clinical outcomes in bladder cancer

**DOI:** 10.1101/2023.02.20.528920

**Authors:** Trine Strandgaard, Iver Nordentoft, Karin Birkenkamp-Demtröder, Liina Salminen, Frederik Prip, Julie Rasmussen, Tine Ginnerup Andreasen, Sia Viborg Lindskrog, Emil Christensen, Philippe Lamy, Michael Knudsen, Torben Steiniche, Jørgen Bjerggaard Jensen, Lars Dyrskjøt

## Abstract

Bladder field cancerization may be associated with disease outcome in patients with bladder cancer. To investigate this, we analyzed biopsies from bladder urothelium and urine samples by genomics and proteomics analyses. Samples were procured from multiple timepoints from 134 patients with early stage bladder cancer and detailed long term follow-up. We measured the field cancerization in normal-appearing bladder biopsies and found that high levels were associated with high tumor mutational burden, high neoantigen load, and high tumor-associated CD8 T-cell exhaustion. Non-synonymous mutations in known bladder cancer driver genes such as *KDM6A* and *TP53* were identified as early disease drivers in normal urothelium. High field cancerization was associated with worse outcome but not with response to BCG. The level of urinary tumor DNA (utDNA) reflected the bladder tumor burden and originated from both tumors and field cancerization. High utDNA levels after BCG were associated with worse clinical outcomes for the patients. Our results indicate that the level of field cancerization may affect clinical outcome, tumor development and immune responses. utDNA measurements have significant prognostic value and reflect the disease status of the bladder.

## Introduction

Epithelial tumors arise from dysplasia or carcinoma *in situ* (CIS) precursor lesions, harboring genomically and histologically altered cells. However, preceding the development of precursor lesions and tumors, the normal-appearing epithelium may contain transformed cells harboring cancer-initiating driver alterations. While previous methods did not provide the required resolution to reveal the mutational landscape of normal-appearing tissues, a high resolution insight into these processes has now become possible using high-throughput deep sequencing approaches.

Field cancerization describes areas of the epithelium affected by expanded clonal mutations, which develop from a cell lineage that acquires genetic mutations. The mutations may result in growth advantages of the clone, positive selection in the microenvironment and subsequent growth leading to development of larger fields. The mutated cells within these fields may be morphologically normal or dysplastic and may predispose to the development of malignancies within the area^1,2^. Field cancerization has been described in bladder cancer (BC) as an explanation of the high recurrence rates and clonal relationships observed between synchronous and metachronous tumors^3–6^. Recent studies performing detailed mapping of the bladder mucosa using high-throughput sequencing approaches have identified widespread field cancerization in the bladder of individuals with and without BC, indicating the necessity to differentiate between fields with malignant potential and fields with mutations inconsequential for BC carcinogenesis^7–9^. Previous studies have suggested that tumor formation may be highly dependent on the genes affected by mutations, the combination of affected genes and the order in which mutations occur^3,10^, being a possible explanation for the tumor development from some but not all fields.

Intravesical treatment with Bacillus Calmette-Guérin (BCG) is part of the standard treatment regimen for patients with high-risk non-muscle invasive BC (NMIBC). The main target of BCG instillation is CIS and field cancerization of the bladder, which remains after surgical resection of the tumors. Whether the extent or the composition of field cancerization has an impact on clinical manifestation, recurrence rates and treatment response, and how the field cancerization may affect the tumor microenvironment (TME), remain to be investigated. The local mutational burden of the bladder, including tumors and field cancerization, as well as renal clearance of the circulating tumor DNA (ctDNA) may be reflected by the presence of urinary tumor DNA (utDNA). Consequently, urinary tests of utDNA may provide a tool for continuous monitoring of the bladder disease state. Previous studies have found high utDNA levels in patients with NMIBC to be associated with worse recurrence-free survival^11^ and progression to muscle invasive disease^12,13^. It is unclear to what extent the bladder field cancerization and renal clearance might impact the utDNA levels. While studies on muscle-invasive bladder cancer have found utDNA levels to be associated with lack of response to neoadjuvant chemotherapy (NAC)^14^, the prognostic and predictive role of utDNA in NMIBC needs further investigation.

Here, we analyzed selected site biopsies (SSBs; n=751) and urine samples (n=187) from multiple bladder locations and clinical visits procured throughout the disease courses of 134 patients with high-risk NMIBC to investigate the prognostic and predictive roles of field cancerization and utDNA.

## Methods

### Patients, follow-up and biological samples

A total of 134 patients with NMIBC were included in this retrospective study. Patients received treatment at Aarhus University Hospital between 1994 and 2018 and provided informed written consent to participate in future research. The study was approved by The Danish National Committees on Health Research Ethics (#1708266). All methods in the study were carried out in accordance with approved guidelines and regulations.

All patients received at least five instillations of BCG. End of follow-up (FU) was defined as the last of the following: last cystoscopy, last detected tumor, cystectomy, progression or metastases. Recurrence-free-survival (RFS) and HG-recurrence-free-survival (HG-RFS) were calculated from the SSB with the highest number of mutations and until first recurrence or HG-recurrence/progression, respectively, or end of FU. Post-BCG HG-RFS was calculated from BCG end-date and until first HG-recurrence or progression to MIBC, or end of FU. Patients were censored at the end of FU in absence of an event. Patients were censored if less than three biopsies were analyzed and no mutations were called (n=2). Post-BCG HG-recurrence status was defined as follows: patients in BCG HG-recurrence group experienced a HG recurrence within two years after end of BCG or progressed to MIBC anytime during their disease course. BCG non-HG-recurrence patients did not develop new HG disease within two years after end of BCG and did not progress to MIBC.

DNA from tumors and leukocytes was analyzed from all 134 patients using whole exome sequencing (WES). Furthermore, SSBs were analyzed from 70 (52%) patients and urine samples from 104 (78%) patients. Normal-appearing SSBs were selected based on the pathologist’s original descriptions of solely normal tissue in the samples. Furthermore, SSBs with atypia, hyperplasia, dysplasia (Grade I and II), carcinoma *in situ* (CIS) and papillary tumor as well as urine samples were analyzed. SSBs were collected primarily before BCG initiation at multiple timepoints during the disease courses. Additionally, for some patients, samples collected after BCG were included. For a detailed overview of included patients and samples, see **Table 1 and Supp. Fig. S1**.

**Table 1.** Clinical characteristics of the selected patient cohort. Samples collected within 120 days after TURBT were considered reTURBT. Analyzed tumors account for the tumor samples (WES or RNA-seq) closest to BCG-initiation or diagnosed tumor. BCG = Bacillus Calmette-Guérin. HG = high grade. LG = low grade/G2. CIS = Carcinoma in situ. SSB = Selected site biopsy. utDNA = urinary tumor DNA. Normal SSBs cover normal-appearing SSBs. Abnormal SSBs include SSBs with tumor, CIS, dysplasia, hyperplasia and metaplasia. TURBT = transurethral resection of bladder tumors. IQR = Interquartile range. *Two samples were missing tumor grade. For simplicity, they have been included as HG tumors.

| Variable | N = 134 |
| --- | --- |
| Age, Median (IQR) | 70 (62, 75) |
| Sex, n (%) |  |
| Female | 29 (22%) |
| Male | 105 (78%) |
| Smoking status, n (%) |  |
| Current | 55 (41%) |
| Former | 60 (45%) |
| Never | 18 (13%) |
| Unknown | 1 (0.7%) |
| Pack years, Median (IQR) | 25 (15, 40) |
| Progression to MIBC, n (%) |  |
| Progression | 28 (21%) |
| No progression | 106 (79%) |
| Total number of tumors, Median (IQR) | 6 (3, 10) |
| Recurrence rate, Median (IQR) | 0.96 (0.52, 1.50) |
| Outcome category, n (%) |  |
| BCG HG-recurrence | 64 (48%) |
| BCG non-HG-recurrence | 70 (52%) |
| Number of normal SSBs sequenced/pt, Median (IQR) | 10 (5, 13) |
| Number of abnormal SSBs sequenced, Median (IQR) | 2.00 (1.00, 3.25) |
| Analyses performed, n (%) |  |
| SSB | 30 (22%) |
| SSB and utDNA | 39 (29%) |
| utDNA | 65 (49%) |
| Pre-BCG stage and grade, n (%) |  |
| CIS | 17 (13%) |
| T1 HG* | 39 (29%) |
| T1 LG | 6 (4.5%) |
| Ta HG* | 38 (28%) |
| Ta LG | 34 (25%) |
| Pre-BCG concomitant CIS, n (%) | 34 (25%) |
| Pre-BCG reTURBT, n (%) |  |
| reTURBT | 32 (24%) |
| No reTURBT | 102 (76%) |
| Pre-BCG tumor status, n (%) |  |
| Primary tumor | 42 (31%) |
| Recurrent tumor | 92 (69%) |
| Pre-BCG EAU risk group, n (%) |  |
| Very High Risk | 29 (22%) |
| High Risk | 53 (40%) |
| Intermediate Risk | 52 (39%) |
| Previous Mitomycin C treatment, n (%) | 8 (6.0%) |
| Post-BCG follow-up (years), Median (IQR) | 6.0 (2.8, 8.4) |
| Follow-up from first analyzed sample (years), Median (IQR) | 7.8 (4.3, 10.0) |

Clinical characteristics of the selected patient cohort. FU = follow-up. HG = High grade. LG = Low grade+G2. CIS = Carcinoma in situ. SSB = Selected site biopsy. tdDNA = Tumor-derived DNA. Normal SSBs cover normal-appearing SSBs. Abnormal SSBs include SSBs with tumor, CIS, dysplasia, hyperplasia and metaplasia.\* Two samples were missing tumor grade. For simplicity, they have been included HG tumors.

Tumor biopsies were either fresh frozen (FF) or from formalin fixed paraffin embedded (FFPE) samples. Blood samples from all patients were stored in EDTA tubes at -80° C. SSBs were provided as FFPE samples. Urine samples were processed as previously described^15^.

### DNA extraction

Tumor DNA from FF and dry-frozen tumors was extracted with Gentra Puregene Tissue Kit (Qiagen). DNA from FFPE samples was extracted using either AllPrep DNA/RNA Kit (Qiagen) for tumors, or GeneRead FFPE Kit (Qiagen) with extra deparaffinization solution (320μl) for SSBs. From peripheral blood, leukocyte DNA for germline (GL) reference was extracted using Qiasymphony DSP DNA midi kit (Qiagen).

Tumor sections (4 μm) were haematoxylin and eosin (HE) stained to estimate carcinoma cell percentage.

DNA from FF and dry-frozen tumors was extracted from 20-25 serial cryosections of 20μm. From FFPE tumor samples and SSBs, DNA was extracted from punches using a 1.5 mm biopsy needle. For SSBs, all material in the block was used for extraction due to very small samples. Total cell-free DNA from urine supernatants was extracted from a median of 3.6 mL urine supernatant (range 0.7–3.65 mL) incubated with 10% ATL buffer before purification using the QIAsymphony DSP Circulating DNA Kit (Qiagen).

### Whole Exome Sequencing and data processing

Next generation sequencing (NGS) libraries and subsequent WES capture were prepared using either RefSeq spike-in probes from Twist Bioscience in combination with Twist Human Core Exome Capture kit or using the illumina TruSeq DNA Kit and NimbleGen SeqCap EZ v3.0. Samples were sequenced using Illumina Sequence platforms. Fastq files were trimmed using cutadapt and mapped with bwa-mem using the GRCh38 genome assembly. Duplicate reads were marked using MarkDuplicates from GATK and base quality scores were recalibrated (ApplyBQSR, GATK). Variants were called using Mutect2 and annotated using SnpEffv4.3i. Finally, variants with frequency below 5% (VAF < 5%), less than three alternate allele reads in the tumor or a ROQ score (Phred-scaled probability that the variant alleles are not due to a read orientation artifact) below 30 were filtered out.

For the estimation of neoantigen load in tumors, HLA types were called using POLYSOLVER^16^, xHLA^17^, and OptiType^18^, with patient HLA type decided by consensus vote supported by at least two algorithms. If no majority could be reached, POLYSOLVER was used. All novel 9-11mer peptide fragments were generated by MuPeXI^19^ and eluted ligands (EL) rank-percentage scores for all HLA alleles were predicted by NetMHCpan-4.1^20^. The rank-percentage score represents the rank of the fragments EL probability compared to a set of random natural peptides. A mutation was considered a neoantigen if at least one fragment had an EL rank percentage score<2%, for at least one HLA allele.

Tumor mutation burden (TMB) was defined as the total number of mutations divided by 36.8 Mb (Size of the Twist Human Core Exome Capture kit panel + Refseq) or 64 Mb (Size of the NimbleGen SeqCap EZ v3.0 panel).

### RNA-sequencing and data processing

RNA-sequencing (RNA-seq) was performed using either ScriptSeq (EpiCentre) library preparation or using the KAPA RNA HyperPrep Kit (RiboErase HMR; Roche) for library preparation. Generated libraries were sequenced on Illumina platforms. Salmon^21^ was used to quantify the expression of transcripts using annotation from the Gencode release 33 on genome assembly GRCh38. Transcript-level estimates were imported and summarized at gene-level using the tximport R library. Samples with less than 5 mill. mapped reads were excluded and genes not expressed in more than 25% of the remaining samples were filtered out.

We estimated immune cell populations from the RNA-seq data using established gene expression signatures as in Rosenthal *et al*.^*22–24*^

### Estimation of CD8 T-cell exhaustion and post-BCG exhaustion predictor (ExhP)

T-cell exhaustion and post-BCG exhaustion predictor (ExhP) were defined as previously described^25^. In short, the residuals from the linear correlation between the estimated level of CD8 T-cells and the mean gene expression level of *PDCD1, CTLA4, LAG3, HAVCR2*, and *KLRG1* were used to estimate the CD8 T-cell adjusted exhaustion level for all tumors. The post-BCG ExhP was defined as the ratio between the identified genes upregulated in pre-BCG tumors from patients having exhausted and non-exhausted tumors post-BCG, respectively. An optimal cutpoint for the post-BCG ExhP according to time to BCG HG-recurrence was used to define the dichotomized level of post-BCG ExhP.

### Design of custom targeted sequencing panels

For high-throughput targeted sequencing of tumor specific mutations, we designed three NGS panels with 10-71 unique mutations from the patients covering 48, 53 and 50 patients respectively, as described in^25^. DNA from SSBs were sequenced using panel 1 and 2, whereas urinary cfDNA was sequenced using all three panels.

Single nucleotide variants (SNVs) for panel inclusion were selected based on the following criteria: (1) high or moderate impact, (2) high variant allele frequencies (VAFs), (3) known oncogenic genes^26^, and (4) bladder cancer associated genes^27,28,29,30,4^. Mutations in the most exonically variable genes (i.e. genes typically poorly sequenced, with high variation in healthy individuals or encoding very long proteins) reported by the Ingenuity Variant Analysis (IVA) software and/or at error-prone positions/in commonly reported erroneous contexts (C>T) were excluded, unless present in cancer driver genes or known bladder cancer genes. The three panels differed slightly in mutation selection. Patient-specific single nucleotide polymorphisms (SNPs) were included on panel 1 for subsequent distinction and quantification of mutations in pooled samples.

### Library preparation for targeted sequencing of DNA from SSBs

DNA from SSBs was sequenced using the designed custom panel for targeted sequencing. Libraries were prepared using the Twist Library preparation EF kit (Twist Bioscience) with an input of 50 ng DNA. The protocol involves enzymatic fragmentation; however, the fragmentation time was decreased to 6 minutes to account for degraded DNA due to aged FFPE blocks and very small biopsies increasing the ratio of formamide contact to tumor tissue and degradation. Prior to library generation of samples sequenced using panel 1, DNA from SSB samples were mixed in pools of 2-8 samples in order to reach sufficient DNA input amounts of 50ng. Samples sequenced using panel 2 were not pooled. Finally, for robust error correction^31^, a 9 bp. Unique Molecular Identifiers (UMI) were incorporated by replacing the twist adaptors with xGen™ UDI-UMI Adapters (Integrated DNA Technologies). Libraries were captured using the Twist Custom Panel described in the previous section. Post-library PCR amplification was set to 8 cycles and post capture amplification to 14 cycles. Sequencing was performed on the NovaSeq 6000 platform (illumina).

### Library preparation for targeted sequencing of utDNA

Libraries and sequencing were performed as described previously^25^. In short, for deep-targeted sequencing of urine supernatants, we used the designed NGS-panels (TWIST Bioscience) and used modified Twist protocols for library preparation and capture in combination with Unique Molecular Identifiers (UMIs).

### Targeted sequencing mutation calling and data processing

Reads were mapped against the hg19 (panel 1) or hg38 (panel 2 and 3) genome using bwa mem v.0.7.17. After mapping, consensus reads were generated from UMIs with at least three identical UMIs supporting each consensus mutation call. Hereafter, read counts for the SNV and SNP positions included in the panel were evaluated using the pileup tool bam-readcount.

For error-robust calling of low-frequency mutations in SSBs and urine samples, we applied the deepSNV pipeline^32,33^.

The inclusion of genomic SNV positions inferred from multiple patients on every custom panel, facilitates an abundance of sequencing data for every genomic position. Positions where no mutations are expected to be present are sequenced for each of the patients due to the patient specific mutations. We exploited this by employing an analysis framework based on a maximum likelihood implementation of the shearwater algorithm developed by Gerstung *et al*.^32^. In brief, a background error model was built by fitting presumably non-mutated data, i.e. data from all samples not associated with a given mutation, to a binomial distribution with site-specific calculation of the dispersion for every mutation of interest. Samples with a VAF at a given position above 10% were excluded when generating the error model. Positions with error rates above 10% were excluded. Presumably mutated positions, i.e. data from the target positions of a given sample, was assessed for a statistical significant difference compared to the background error model. A rho value of 10^−4^ was used and only the specific base changes selected in the design of the panel were considered for mutational analysis of the relevant genomic positions. Five SNPs were included for each patient in order to adjust VAF for samples being part of a pool. Resulting *p*-values were corrected for multiple testing using the Benjamini-Hochberg procedure and adjusted *p*-values below 0.05 were considered significant.

For analysis of driver mutations in KEGG pathways, gene lists representing pathways were obtained using R package OmnipathR for PI3K-Akt signaling and p53 signaling, whereas gene lists for chromosome related pathways were obtained from https://www.genome.jp/brite/hsa03036+1105. Genes not related to these, were classified as “Other”, and pathways were prioritized in the following order when genes were represented in multiple pathways: p53 signaling, PI3K-Akt signaling, Chromosome, Other.

### Field cancerization measure

The degree of field cancerization was estimated as follows: the number of mutations in the most mutated sample from a patient (before, after or across the whole disease course) adjusted for each patient’s mutational weight on the panel. This was done by multiplying the fraction of positions on the panel for each patient out of the maximum number of mutations on the panel for any patient. For every patient, the following was calculated: 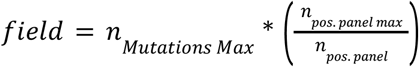 (**Fig. 1a**).

**Fig. 1.**
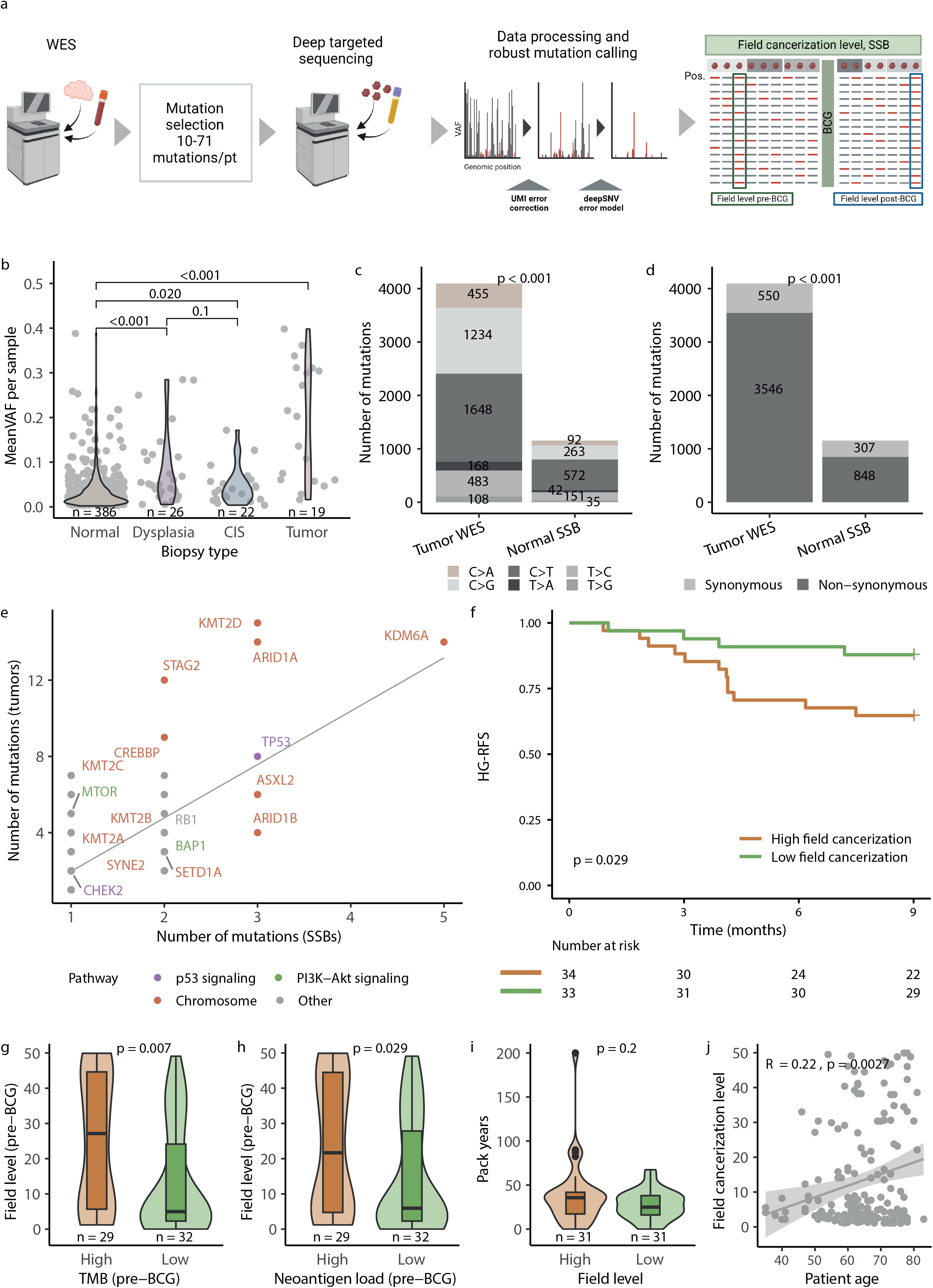
Study overview and analyses of selected site biopsies (SSBs). a) Overview of the process of data acquisition including panel design from tumor WES data for deep targeted sequencing of DNA from SSBs and urine samples, data processing and estimation of field cancerization level from SSBs (created with BioRender.com). b) Mean VAF measured per sample across different sample types analyzed with deep targeted sequencing of SSB DNA (Wilcoxon rank sum test and Bonferroni adjustment). c) Analysis of the number of mutations of six alteration types in tumors and normal appearing SSBs (Fisher’s exact test). d) Analysis of the number of synonymous (low+modifier impact) and non-synonymous (high +moderate impact) mutations in tumors and normal appearing SSBs (Fisher’s exact test). e) Number of non-synonymous mutations in normal appearing SSBs and tumor samples in cancer driver genes colored by functional pathways for genes with more than five mutations. f) Kaplan-Meier plot of high grade recurrence free survival (HG-RFS) from SSBs with highest number of mutations for 67 patients stratified by level of field cancerization level. Patients without any detectable field and less than three analyzed SSBs were left out from the analysis. Field groups were based on median (log-rank test). g) Comparison of the pre-BCG field cancerization level and the pre-BCG tumor mutation burden, split by median (Wilcoxon rank sum test). h) Comparison of the pre-BCG field cancerization level and the pre-BCG tumor neoantigen load (Wilcoxon rank sum test). i) Comparison of the field cancerization level, median split, and the number of pack years for current or former smokers (Wilcoxon rank sum test). j) Correlation between the patient age at sampling time and the level of field cancerization in the all analyzed samples with measurable mutations (Pearson correlation).

### Statistical analysis

For categorical variables, Fisher’s Exact test was used. Continuous data was tested using Wilcoxon Rank Sum test (unpaired data). Categorical variables were compared using Fisher’s exact test. For multiple testing, Bonferroni correction was applied. Kaplan-Meier curves with associated log-rank tests were performed to assess time to HG-RFS, post-BCG HG-RFS or RFS. Pearson correlation was used for comparison of numerical values. Differences in protein expression in urine samples between relevant groups were measured using unpaired t-tests. Statistical significance was set at p<0.05, except for protein expression analyses in Volcano plots, where it was set at q<0.1. All statistical analyses were performed using R version 4.1.1.

## Results

### Patient characteristics and molecular analyses

We analyzed samples from a total of 134 patients with NMIBC that received at least five BCG instillations. The patients were followed for a median of 7.8 years after the first analyzed sample and 6.0 years after BCG. Samples were analyzed as illustrated in **Fig. 1a and Supp. Fig. S1**. Samples were procured at multiple timepoints and bladder locations throughout the patient disease courses. See **Table 1** for clinical information.

The level of field cancerization was defined as the maximum number of mutations detected at any sample collected anytime during the disease course (considering both pre- and post-BCG samples) adjusted for the patient’s mutational weight on the panel (see methods and **Fig. 1a**).

### Field cancerization is associated with tumor biology

We performed deep targeted sequencing of DNA from 751 SSBs collected throughout 70 patients’ disease courses. This included 662 biopsies of normal appearing urothelium, 79 dysplastic/cancerous lesions and 10 SSBs with other abnormal characteristics. Samples were sequenced to a mean coverage of 19,338X before UMI-consolidation and 1,357X after UMI-collapsing. We detected tumor specific mutations in 458 out of 751 analyzed samples. In mutated samples, we detected a mean of 17% of the mutations included on the panel for the patient in question.

We compared the mean variant allele frequency (VAF) per sample between different sample types and observed that mutations found in normal appearing SSBs had low variant allele frequencies (mean=0.036). The VAF per sample was observed to increase from normal-appearing urothelium, across dysplastic lesions to tumor samples (**Fig. 1b**), indicating an enrichment of transformed cells in the specimens. To further explore the role of field cancerization in tumor development, we analyzed the mutational characteristics of tumor mutations (included on the panels for deep targeted sequencing) in normal appearing SSBs. We found that although a subset of the tumor mutations were detected in normal appearing biopsies, the number of C>T mutations constitute a large fraction of mutations in SSBs (**Fig. 1c**). C>T mutations have previously been described in normal bladder tissue and accumulate in normal cells with age^7,9,34^. Additionally, we observed that non-synonymous mutations were more relatively abundant in tumor samples compared to normal appearing SSBs, where synonymous mutations were more often detected (**Fig. 1d**). We further investigated whether specific cancer driver genes were mutated in normal SSBs, potentially indicating a cancer-driver in the field, and found that 33% of high impact mutations in cancer driver genes originally observed in the tumor samples were already present in normal SSBs (**Table 2**). Among the cancer driver genes found to be already mutated in normal SSBs were *TP53, STAG2, ARID1A, KMT2D* and *PIK3CA* (**Fig. 1e**), known to be important genes in BC development and progression^35^.

**Table 2.** Analysis of cancer driver genes in tumors and normal SSBs. The full list of high impact mutations in cancer driver genes detected in tumor and normal-appearing SSBs as well as only in tumor samples.

| Driver gene | Tumor WES | Normal SSB | Driver gene | Tumor WES | Normal SSB | Driver gene | Tumor WES | Normal SSB |
| --- | --- | --- | --- | --- | --- | --- | --- | --- |
| *EGFR* | Mutation | Mutation | *SETDB2* | Mutation | No mutation | *SPTAN1* | Mutation | No mutation |
| *NCOR1* | Mutation | Mutation | *BUB1B* | Mutation | No mutation | *CYP17A1* | Mutation | No mutation |
| *U2AF1* | Mutation | Mutation | *CLSPN* | Mutation | No mutation | *MIB1* | Mutation | No mutation |
| *PREX2* | Mutation | Mutation | *EP300* | Mutation | No mutation | *MAPK1* | Mutation | No mutation |
| *STAG2* | Mutation | Mutation | *KIAA1109* | Mutation | No mutation | *DDR2* | Mutation | No mutation |
| *RB1* | Mutation | Mutation | *CDKN1A* | Mutation | No mutation | *STAT6* | Mutation | No mutation |
| *LYST* | Mutation | Mutation | *ATG5* | Mutation | No mutation | *HDAC9* | Mutation | No mutation |
| *ARID1A* | Mutation | Mutation | *RAD51B* | Mutation | No mutation | *IRF1* | Mutation | No mutation |
| *KMT2D* | Mutation | Mutation | *POLE* | Mutation | No mutation | *RUNX1* | Mutation | No mutation |
| *CHEK2* | Mutation | Mutation | *MEN1* | Mutation | No mutation | *TOP2A* | Mutation | No mutation |
| *FBXW7* | Mutation | Mutation | *TRIM27* | Mutation | No mutation | *DDX10* | Mutation | No mutation |
| *IKZF2* | Mutation | Mutation | *ARID2* | Mutation | No mutation | *PRKDC* | Mutation | No mutation |
| *CDC20* | Mutation | Mutation | *DDX4* | Mutation | No mutation | *ZMYM3* | Mutation | No mutation |
| *SP140* | Mutation | Mutation | *LRP1B* | Mutation | No mutation | *CDKN2A* | Mutation | No mutation |
| *DDX3X* | Mutation | Mutation | *PTPN6* | Mutation | No mutation | *RTEL1* | Mutation | No mutation |
| *TP53* | Mutation | Mutation | *CSF3R* | Mutation | No mutation | *CD44* | Mutation | No mutation |
| *COL3A1* | Mutation | Mutation | *HECTD1* | Mutation | No mutation | *RYBP* | Mutation | No mutation |
| *ASXL2* | Mutation | Mutation | *CRKL* | Mutation | No mutation | *CSF1R* | Mutation | No mutation |
| *PSIP1* | Mutation | Mutation | *GRHL3* | Mutation | No mutation | *RARA* | Mutation | No mutation |
| *RANBP2* | Mutation | Mutation | *POLQ* | Mutation | No mutation | *SOS1* | Mutation | No mutation |
| *ATP2B3* | Mutation | Mutation | *TSC1* | Mutation | No mutation | *PGR* | Mutation | No mutation |
| *KDM6A* | Mutation | Mutation | *NFATC2* | Mutation | No mutation | *BRAF* | Mutation | No mutation |
| *KMT2A* | Mutation | Mutation | *HRAS* | Mutation | No mutation | *BAP1* | Mutation | No mutation |
| *TRIM29* | Mutation | Mutation | *PC* | Mutation | No mutation | *BCL9L* | Mutation | No mutation |
| *PTPRB* | Mutation | Mutation | *RBL2* | Mutation | No mutation | *AXL* | Mutation | No mutation |
| *CREBBP* | Mutation | Mutation | *SAMHD1* | Mutation | No mutation | *PTPRD* | Mutation | No mutation |
| *SPOP* | Mutation | Mutation | *DIS3* | Mutation | No mutation | *RNF213* | Mutation | No mutation |
| *PIK3R1* | Mutation | Mutation | *EBF1* | Mutation | No mutation | *DDB2* | Mutation | No mutation |
| *SYNE2* | Mutation | Mutation | *BUB1* | Mutation | No mutation | *ATRX* | Mutation | No mutation |
| *DOT1L* | Mutation | Mutation | *KMT2B* | Mutation | No mutation | *SMARCA4* | Mutation | No mutation |
| *PIK3CA* | Mutation | Mutation | *DDR1* | Mutation | No mutation | *RHOB* | Mutation | No mutation |
| *PTPN13* | Mutation | Mutation | *HUWE1* | Mutation | No mutation | *RANBP17* | Mutation | No mutation |
| *ESCO2* | Mutation | Mutation | *AFF1* | Mutation | No mutation | *FH* | Mutation | No mutation |
| *WRN* | Mutation | Mutation | *ZNF750* | Mutation | No mutation | *AMER1* | Mutation | No mutation |
| *ZFHX3* | Mutation | Mutation | *AKAP9* | Mutation | No mutation | *MAD2L2* | Mutation | No mutation |
| *MDN1* | Mutation | Mutation | *ATM* | Mutation | No mutation | *VPS13C* | Mutation | No mutation |
| *PDE4DIP* | Mutation | Mutation | *FANCD2* | Mutation | No mutation | *PARP1* | Mutation | No mutation |
| *KMT2C* | Mutation | Mutation | *SMARCA2* | Mutation | No mutation | *MAPK3* | Mutation | No mutation |
| *RBM10* | Mutation | Mutation | *RALGDS* | Mutation | No mutation | *FGFR2* | Mutation | No mutation |
|  |  |  | *ARHGEF12* | Mutation | No mutation | *YES1* | Mutation | No mutation |

A high field cancerization level was associated with worse high-grade recurrence-free survival (HG-RFS; p=0.029) evaluated shortly after detection (time span between two cystoscopy follow-up visits; **Fig. 1f** and **Supp. Fig. S2a)**, but no significant associations to recurrence-free and progression-free survival were observed (**Supp. Fig. S2b-c**).

A high tumor mutational burden (TMB) and neoantigen load of the last tumor analyzed before BCG was associated with a high pre-BCG level of field cancerization (*p*=0.007 and *p*=0.029; **Fig. 1g-h**). Too few samples were included in the post-BCG setting to perform robust statistical analysis. Surprisingly, smoking was not associated with field manifestation, as no statistically significant correlations between the number of pack years and level of field cancerization were observed (*p*=0.2; **Fig. 1i**). The level of field cancerization increased significantly with higher age (*p*=0.0027; **Fig. 1j**).

To characterize the mutational landscape across the disease course and in different sample types, we selected three patients with multiple different sample types analyzed. The detected VAF at every genomic position was analyzed in the different sample types. Mutations were detected across different sample types indicating a clonal cellular origin (**Supp. Fig. S3a**,**c**,**e**). However, a high level of inter- and intrapatient heterogeneity was observed. Mutations detected in tumor samples had higher VAF compared to the other sample types, regardless of sequencing method (WES or deep targeted sequencing; **Supp. Fig. S3a**,**c**,**e**). Detailed analysis of tumor (WES) and normal-appearing SSBs from the three patients revealed high inter- and intrapatient heterogeneity in terms of number of mutations and VAF of mutations detected in samples collected throughout the patients’ disease courses and at different locations in the bladder (**Supp. Fig. S3b**,**d**,**f**).

### Urinary tumor DNA levels are prognostic and reflect bladder disease status

In addition to analyzing field cancerization by sequencing of SSBs, we performed deep targeted sequencing of 187 utDNA samples procured from 104 patients before and after BCG. This corresponds to a measure of release of cellular DNA from tumor cells and field cancerization as well as from renal clearance of ctDNA. Samples were sequenced to 2,153x UMI consolidated coverage (18,856X raw coverage). Detailed clinical follow-up allowed us to compare utDNA levels with the clinical status of the bladder at multiple clinical time points. The level of utDNA was significantly higher in the presence of tumors at the time of urine sampling. The levels of utDNA increased with higher tumor stage (**Fig. 2a**) and tumor multiplicity (**Fig. 2b**). Interestingly, utDNA was also detected in several cases where no tumor was diagnosed (**Fig. 2a-b**). We therefore analyzed the prognostic importance of this finding and observed a difference in post-BCG RFS between patients with and without detectable utDNA after BCG (*p*=0.072; **Fig. 2c**), potentially indicating a positive lead time between utDNA detection and recurrence development. Additionally, we found that patients progressing to MIBC after BCG had higher levels of utDNA before and after BCG (*p*=0.052 and *p*=0.0014, respectively; **Fig. 2d**). Patients with high recurrence rates (one or more tumors per year) had high utDNA levels after BCG (*p*<0.001; **Fig. 2e**). No differences were observed between patients with high and low pre-BCG utDNA levels and their post-BCG HG-RFS (p=0.18; **Supp. Fig. S2d**). Analysis of utDNA levels after BCG revealed significantly worse post-BCG HG-RFS in patients with high levels of utDNA, indicating treatment failure (*p*=0.047; **Fig 2f**).

**Fig. 2.**
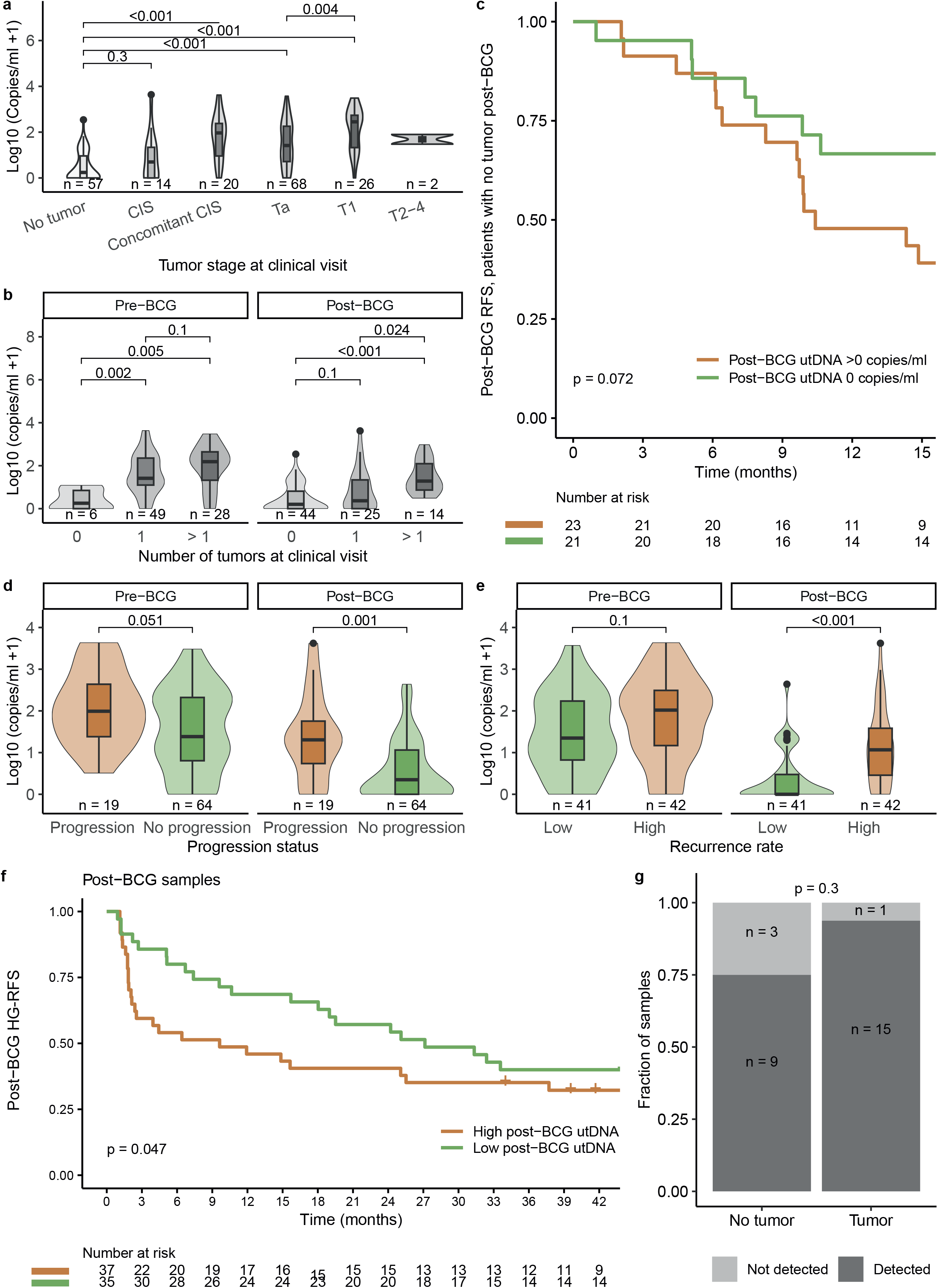
Urinary tumor DNA (utDNA) and clinical correlations. a) Comparison between the utDNA level and tumor status of the bladder at the same clinical time point (Wilcoxon rank sum test, Bonferroni correction). b) Comparison of the number of tumors in the bladder and utDNA level measured at the same clinical visit (Wilcoxon rank sum test). c) Kaplan-Meier plot of post-BCG recurrence-free survival (RFS) for 44 patients where no tumor was clinically visible, but utDNA was measured or absent (log-rank test). d) Comparison of the level of urinary utDNA measured pre- and post-BCG and the progression status of the patients (Wilcoxon rank sum test). e) Comparison of tumor recurrence rate during the disease course and the level of utDNA measured pre- and post-BCG, respectively (High = one or more tumors per year. Low = less than one tumor per year; Wilcoxon rank sum test). f) Kaplan-Meier plot of post-BCG high grade recurrence free survival (HG-RFS) for 72 patients stratified by the post-BCG level of urinary utDNA (split by median; log-rank test). g) Comparison of the fraction of visits where mutations detected in normal appearing SSBs were observed in urine samples from the same clinical visit. Clinical visits with and without tumor in the bladder are shown (Fisher’s exact test). CIS = Carcinoma in situ. P=Progression to MIBC after BCG. NP= No progression.

To compare utDNA mutations and levels to field cancerization status, we compared mutations observed in SSBs to mutations in urinary DNA from samples collected at the same clinical visits (n_visits_ = 28; n_patients_=23) with and without the presence of bladder tumors. Mutations observed in SSB samples were observed in 9/12 urine samples from the same visits when no tumor was found in the bladder, indicating contribution from the field or undetected tumor to the total utDNA level. If a tumor was present at the time of sampling, 15/16 urine samples contained mutations also observed in SSBs. This corroborates that utDNA can originate from the field (or undetected tumors), and not just from detected tumors (*p*=0.3; **Fig. 2g**).

### Field cancerization and utDNA levels may be associated with tumor immunology and BCG treatment response

Previously, we estimated the level of CD8 T-cell exhaustion in pre- and post-BCG tumors from the patients based on gene expression levels of selected immune-inhibitory genes adjusted for the estimated CD8 T-cell infiltration^25^. Furthermore, we developed a predictor of post-BCG CD8 T-cell exhaustion based on pre-BCG tumor gene expression levels. We found that CD8 T-cell exhaustion as well as prediction of post-BCG CD8 T-cell exhaustion were associated with HG recurrences after BCG, indicating treatment failure^25^. Here, we correlated these measures to the level of field cancerization and found high CD8 T-cell exhaustion in pre-BCG tumors to be associated with high pre-BCG field cancerization level (*p*=0.017; **Fig. 3a**). There was no correlation between the pre-BCG CD8 T-cell exhaustion level and the level of post-BCG field cancerization (*p*=0.9; **Supp. Fig. S2e**). Additionally, patients predicted to have high post-BCG exhaustion had high field cancerization levels (*p*=0.065; **Fig. 3b**). There was no association between the level of field cancerization and the amount of CD8 T-cell infiltration in tumors based on deconvolution of RNA-sequencing data (*p*=0.10; **Fig. 3c**). As previously reported, higher levels of utDNA were observed when tumor-infiltrating CD8 T-cells were categorized as exhausted using our CD8 T-cell adjusted exhaustion score^25^ compared to tumor samples with lower CD8 T-cell exhaustion both pre- and post-BCG (*p*=0.008 and *p*= 0.012, respectively; **Fig. 3d**). In the current study, we investigated the correlations between the level of utDNA and the post-BCG exhaustion prediction (ExhP) score which is based on gene expression levels in pre-BCG tumors. It was observed that high ExhP was associated with high pre- and post-BCG levels of utDNA (*p*=0.002 and *p*=0.040, respectively; **Fig. 3e**). The pre-BCG field cancerization level was neither associated with post-BCG HG-recurrence status (*p*=0.3; **Fig. 3f**) nor with post-BCG HG-RFS (*p*=0.93; **Fig. 3g**), suggesting a potential indirect effect of field cancerization on BCG treatment response via effects on the tumor biology. However, field cancerization did not show predictive value per se.

**Fig. 3.**
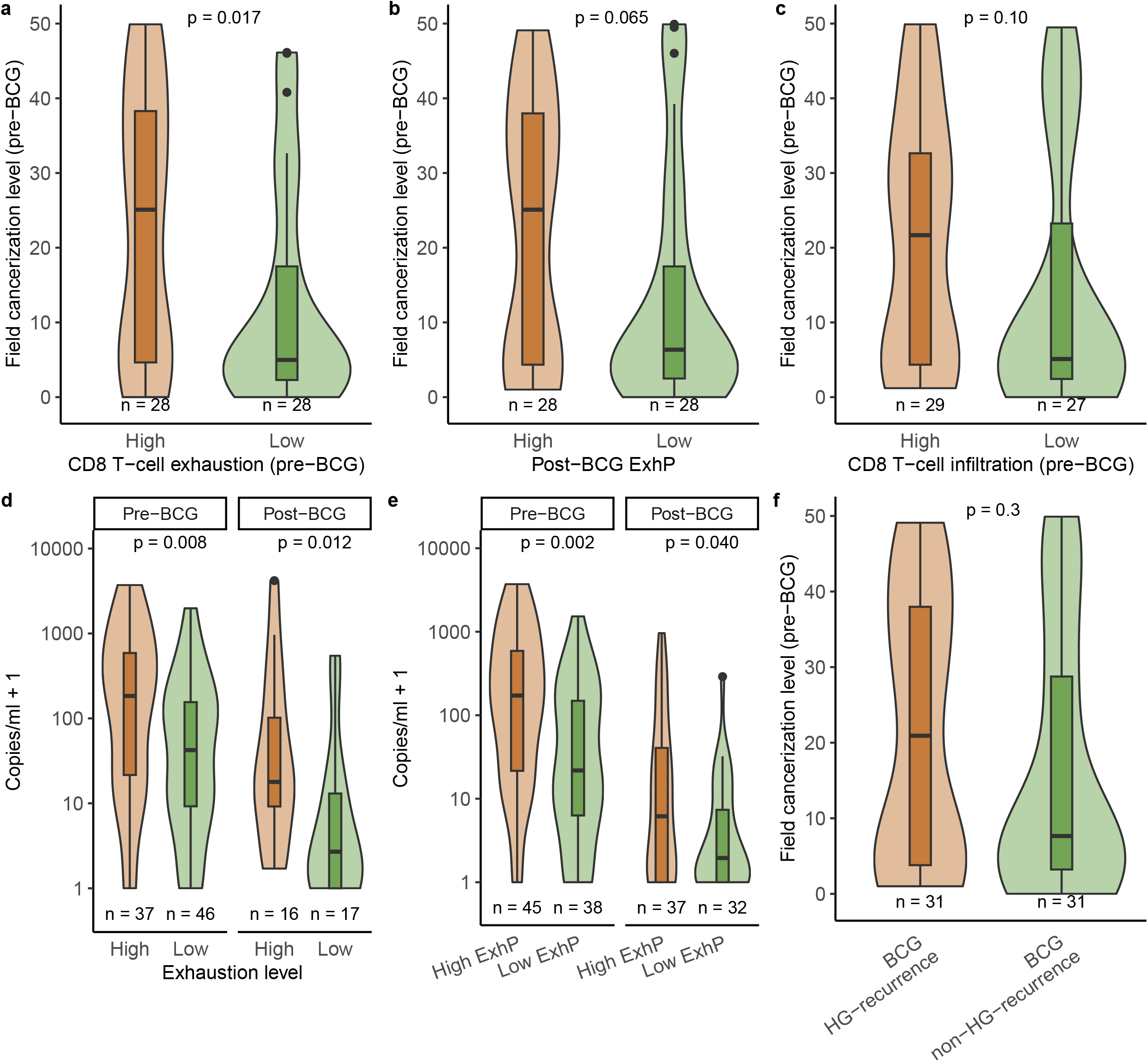
Field cancerization level, utDNA and immuno-related parameters. a-e: Immune correlates estimated from RNA-sequencing data from tumors. a) Comparison of the pre-field cancerization level and the CD8 T-cell adjusted exhaustion status of pre-BCG tumors (Wilcoxon rank sum test). b) Comparison of the pre-field cancerization level and the pre-BCG tumor-based predictor of post-BCG CD8 T-cell exhaustion (ExhP; Wilcoxon rank sum test). c) Comparison of the pre-field cancerization level and the level of CD8 T-cell infiltration in pre-BCG tumors (Wilcoxon rank sum test). d) Comparison of the CD8 T-cell adjusted exhaustion status of pre- and post-BCG tumor samples, respectively, and the level of utDNA (Wilcoxon rank sum test). e) Comparison of the post-BCG exhaustion predictor (ExhP) and the level of utDNA based on pre- and post-BCG measurements, respectively (Wilcoxon rank sum test). f) Comparison of the level of pre-BCG field cancerization level and post-BCG HG-recurrence status of the patients (Wilcoxon rank sum test).

### Field cancerization and utDNA levels reflect urinary immune oncology-related proteins associated with disease aggressiveness

We investigated if the level of field cancerization was reflected in urinary protein levels. We analyzed 91 urine samples from 53 patients for the levels of proteins related to immuno-oncology pathways. Only urine samples collected at clinical visits without detectable tumors were included. We observed that patients with high post-BCG field cancerization levels had significantly higher levels of the proteins VEGFA, CD27, LAP TGFβ1 and TRAIL, amongst others (*p*<0.05; **Fig. 4a**). These proteins have been associated with increased angiogenesis, cell survival, proliferation and migration as well as immune activation^36–38^. Pre-BCG samples were not included due to low numbers when adjusting for tumor status. For many of the patients, the same urine samples were analyzed for both immuno-oncology-related protein expression and utDNA (83/86 pre-BCG urine samples and 69/69 post-BCG urine samples). When comparing the urinary protein levels in post-BCG samples from visits without detectable tumor, IL8 was found to be expressed at significantly higher levels in samples from patients with high utDNA levels compared to low utDNA levels (**Fig. 4b**). IL-8 is a mediator of inflammatory responses and is associated with migration, invasion, angiogenesis and metastasis in cancer^39^. No samples from the pre-BCG setting had high levels of utDNA when no tumor was present in the bladder.

**Fig. 4.**
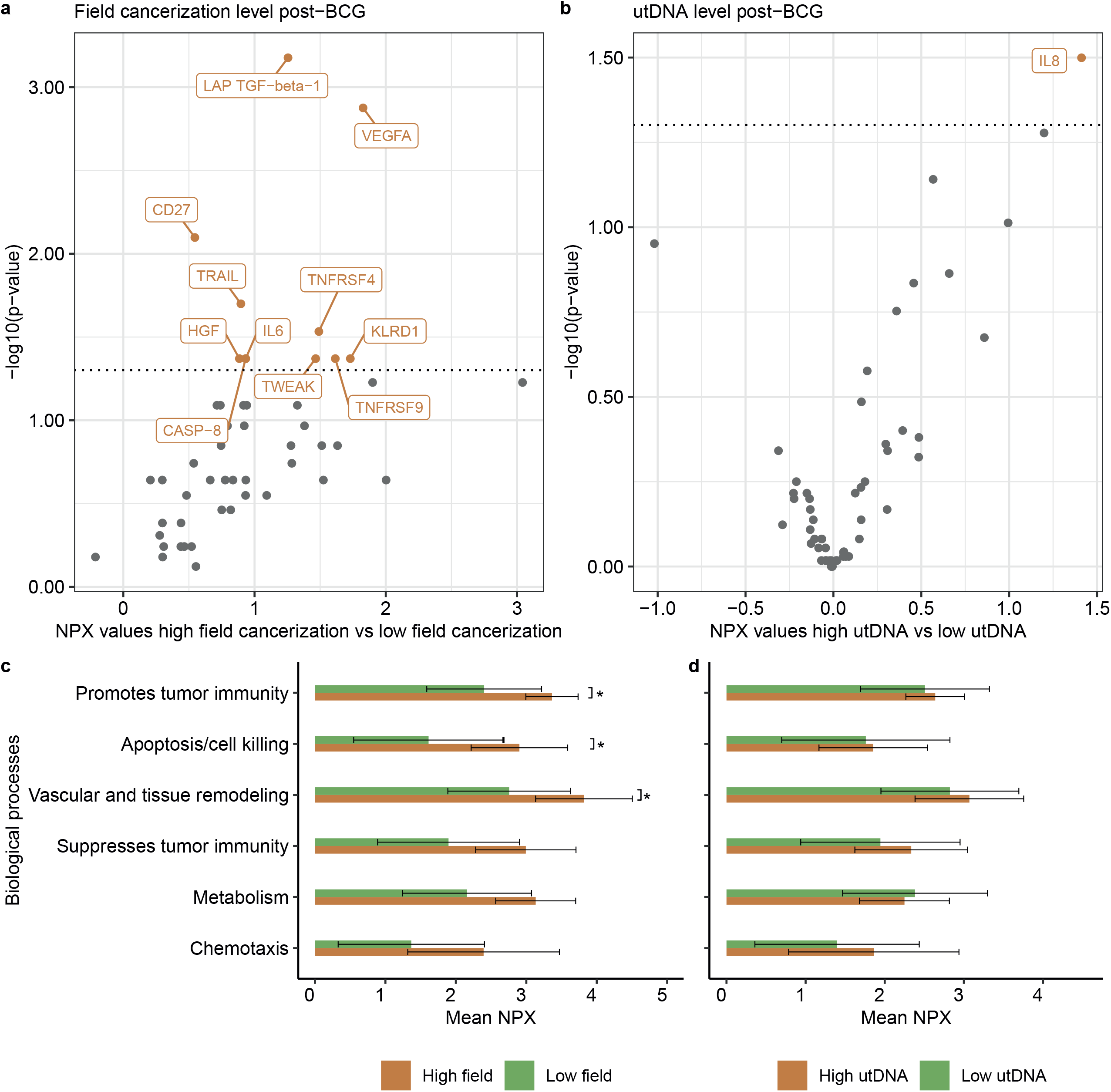
Field cancerization, utDNA and immuno-oncological protein features. a-d) Detection of immune-oncology related protein responses by urine analyses (Olink proteomics). All significantly differentially expressed proteins are named (Wilcoxon Rank Sum test). Colored = significant p-value < 0.05 (unadjusted). Grey = not significant. Dotted lines indicate a significance level of 0.05. Numerical differences in mean NPX-values between compared groups are shown on the x-axis. a) Comparisons between urinary protein levels in patients with high and low field cancerization (split by median) measured in urine samples collected post-BCG (n_patients_=14). b) Comparisons between urinary protein levels in patients with high and low utDNA levels (split by median) measured in urine samples collected post-BCG (n_patients_=36). c) Comparison of the mean urinary level of proteins grouped by biological features between samples from patients with high and low field cancerization post-BCG (n=14; Wilcoxon rank sum test). Non-adjusted p-values are indicated. d) Comparison of the mean urinary level of proteins grouped by biological features between samples from patients with high and low utDNA in post-BCG urine samples (n=36; Wilcoxon rank sum test). Non-adjusted p-values are indicated.

When analyzing the levels of proteins grouped by their biological functions, higher levels of proteins related to suppression of tumor immunity, vascular and tissue remodeling and apoptosis/cell killing were observed after BCG in patients with a high field cancerization level (**Fig. 4c**). For utDNA, no statistically significant differences were observed when protein levels and utDNA were assessed in urine samples collected after BCG treatment (**Fig. 4d**).

## Discussion

Field cancerization has been investigated during several years in multiple cancer types, however, clinical consequences have not been addressed in detail earlier due to small cohort sizes and limited granularity in measurements.

Using deep targeted sequencing, we analyzed 751 normal-appearing and dysplastic/cancerous SSBs along with utDNA from 134 patients with NMIBC.

We found high field cancerization to be associated with HG recurrences, high TMB, tumor neoantigen load and CD8 T-cell exhaustion in tumors as well as with prediction of high post-BCG CD8 T-cell exhaustion, and patient age (**Fig. 5**). In addition, we observed that the levels of utDNA correlated with disease aggressiveness, tumor multiplicity and were associated with tumor CD8 T-cell exhaustion and prediction of post-BCG CD8 T-cell exhaustion, possibly reflecting a more aggressive disease. Interestingly, field cancerization and/or potentially undetected tumors were reflected in utDNA measurements.

**Fig. 5.**
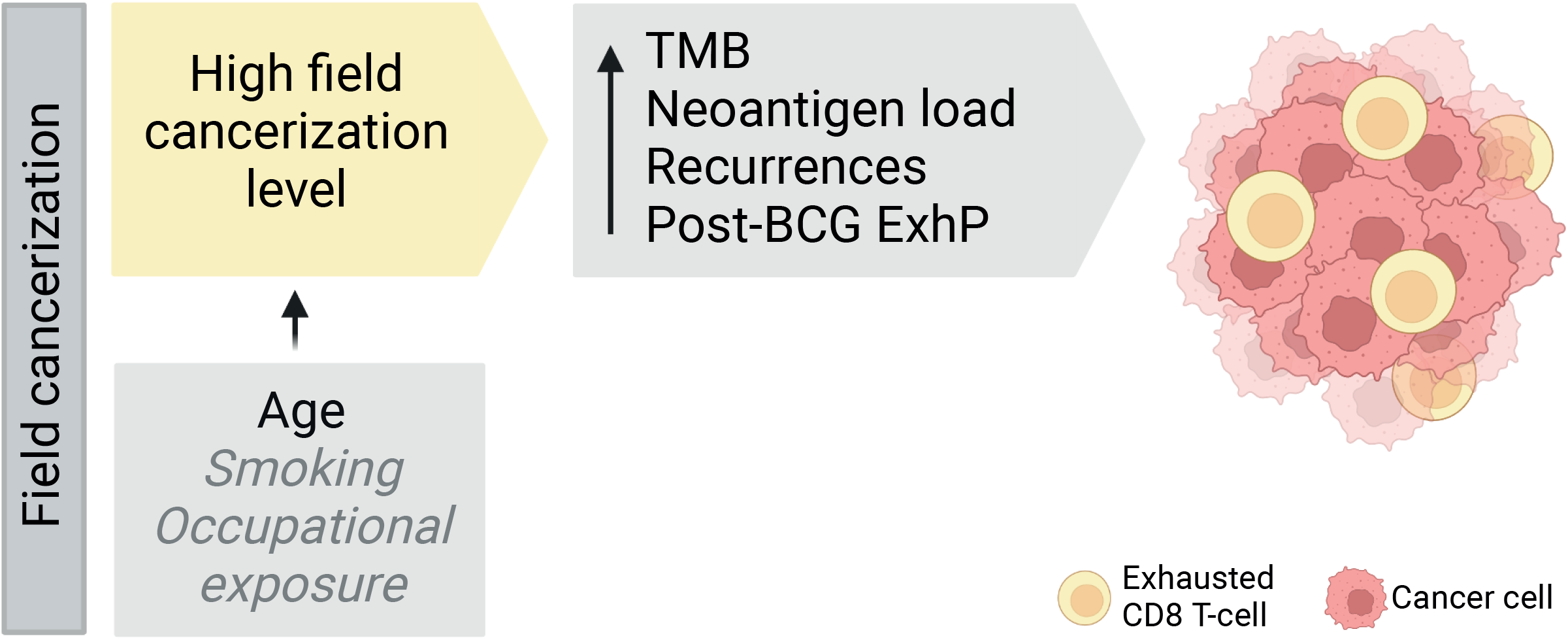
Graphical illustration of summary of main study findings from analyses of field cancerization (created with BioRender.com). Light gray, italic text indicates proposed hypotheses. TMB = Tumor mutation burden. BCG = Bacillus-Calmette Guérin. ExhP = Exhaustion predictor.

We found that high pre-BCG field cancerization levels were associated with high CD8 T-cell exhaustion in BCG-naïve tumors and with prediction of post-BCG exhaustion. Contrarily, we did not observe any differences in the number of tumor-infiltrating CD8 T-cells based on field cancerization levels. Previous studies have shown that prolonged exposure to antigens, either viral or cancerous, may cause CD8 T-cells to become exhausted, resulting in altered effector functions and high expression of immune-inhibitory receptors^40^. We suggest that increased field cancerization may result in prolonged exposure due to continuous presentation of neoantigens, causing CD8 T-cell exhaustion. Abdel-Hakeem *et al*. have shown that previously exhausted T-cells develop into a compromised memory CD8 T-cell phenotype as compared to traditional memory T-cells, resulting in constrained future immune responses^41^. This finding may explain the relationship between high field cancerization levels and CD8 T-cell exhaustion within tumors and prediction of post-BCG exhaustion, although the number of CD8 T-cells were similar. Additionally, we showed that an increased field cancerization level was associated with higher TMB and neoantigen load in the tumors, indicating increased mutagenesis and possible presentation of neoantigens, potentially resulting in exhausted CD8 T-cells. However, additional studies analyzing the immune landscape related to field cancerization are needed to elucidate to interaction between the two.

Our mutational analyses showed that non-synonymous mutations in known cancer driver genes were present in normal appearing SSBs. These genes may represent early tumor drivers present in the normal appearing urothelium. Driver genes only found to be mutated in tumor samples were generally less common in BC and may represent later events in the tumor evolution, although the study design does not allow us to determine tumor specificity. We found mutations to be shared at varying frequencies between various sample types collected at different clinical timepoints. We further found that the level of field cancerization was not predictive of response to BCG-therapy. However, our study showed that cancerous lesions in the bladder and the normal-appearing urothelium share (driver) mutations, indicating a clonal origin from a pre-existing field - or shedding/migration of cancer cells, which is also a possibility^42^. However, this may be more unlikely because of the need for attachment and subsequent growth from the urothelium^6^. Bondaruk *et al*. performed detailed genomic mapping of cancerous bladders and showed that oncogenic mutations were present in normal-appearing and low-grade dysplastic urothelial samples and that tumors developed from field effects in the bladder^8^, supporting the presented findings. The diffuse distribution of mutations at varying frequencies, either because of possible diffuse field cancerization, lack of representative biopsies or selection difficulties with clonal mutations for the sequencing panel design, may complicate the use of SSBs for prognostic and predictive purposes, however, the analyses highlight important aspects of bladder cancer recurrence patterns and tumor driver events.

Consistent with what has been reported by Lawson *et al*.^*7*^, we found field cancerization to increase significantly with age and being characterized by frequent C>T mutations, supporting the accumulation of mutations during life. A high level of field cancerization was associated with worse HG-RFS shortly after the biopsy timepoint, indicating increased risk of development of recurrences with a more altered urothelium. We were not able to show a significant correlation between smoking history (measured in pack years) and the level of field cancerization, although there were indications of higher field cancerization level in patients with more pack years. Smoking is the main risk factor for developing BC, possibly through increased field cancerization induced by smoking carcinogens. A study on field cancerization in lung tissue found that mutation rates of former smokers were similar to never smokers, suggesting a normalization of the field level upon smoking cessation^43^. However, similar studies have not been conducted in bladder specimens. This finding in lung tissue may, at least partly, explain the lack of correlation between smoking exposure and field cancerization level. Furthermore, uncertainties regarding occupational exposure as well as other potential correlates to field cancerization may complicate interpretation of this analysis. Therefore, larger studies with more detailed information on smoking and occupational exposure need to be performed to determine the role of these factors on the development and manifestation of field cancerization and subsequent tumor development and recurrences.

We observed that mutations observed in SSBs were detected in the urine in 9/12 cases when no tumor was found in the bladder. This indicates that the field cancerization may contribute to the utDNA via shedding of mutated DNA. This corroborates the finding in **Fig. 2B-C**, where utDNA was detected without tumors being present in the bladder. However, combined with the finding that utDNA levels increase from visits with only normal urothelium to visits with tumor suggests that the main contributing factor to utDNA may be shedding from existing tumors and not the field cancerization itself. We also observed that higher utDNA levels were correlated to tumor multiplicity, higher recurrence rates and progression to MIBC. Additionally, we found that high post-BCG levels of utDNA, indicating treatment failure, were associated with worse post-BCG HG-RFS. Combined, these results suggest that utDNA may serve as an alternative approach for disease surveillance in BC and treatment response monitoring, mirroring the total mutational status of the bladder by reflecting both field cancerization, multiplicity of tumors and tumor stage as well as risk of post-BCG recurrences. Higher levels of utDNA with higher grade, T-stage and presence of CIS have been reported by Zhang *et al*.^*11*^. Furthermore, others have described utDNA to predict recurrences, treatment failure and progression, suggesting the use of utDNA in the surveillance and prognostic setting^11,12,14,44^. Here we present a detailed, multi omics study on a high number of patients. For clinical implementation the findings need to be validated in clinical trials.

Finally, we investigated the levels of immuno-oncological proteins in the urine from patients with high vs. low field cancerization and high vs. low utDNA levels. Interestingly, we observed that a high field remaining after BCG therapy was associated with higher levels of the proteins VEGFA, CD27 and TRAIL, amongst others. These proteins have been associated with increased vascular formation, cancer cell survival, proliferation and migration as well as immune activation^36–38^, suggesting an altered, immune modulating and cancer-promoting urothelium when affected by high field cancerization. High utDNA levels after BCG were associated with increased levels of the mediator of inflammatory responses, IL-8, suggesting that the presence of field cancerization, as indicated by the shedding of mutated DNA to the urine in the absence of tumors, may result in immune activation in the bladder.

Altogether, these results suggest that field cancerization forms the basis of clonal development of dysplastic and cancer lesions in the bladder and may be important for driving an exhausted phenotype of CD8 T-cells. Furthermore, utDNA may serve as a prognostic and predictive biomarker in BC, potentially improving the clinical management of patients with NMIBC.

## Conclusions

Our results show several important correlates to BC carcinogenesis, prognosis and treatment outcome. High levels of field cancerization in the bladder may cause CD8 T-cell exhaustion in tumors, possibly via increased TMB and neoantigen loads within tumors. Furthermore, high field cancerization is associated with worse clinical outcomes for the patients. The overall mutational status of the bladder, including field cancerization, is reflected in urine samples and utDNA measurements may be used for assessing risk in NMIBC, and for guiding surveillance and early treatment interventions. The findings need to be validated in clinical trials, but could eventually improve surveillance and treatment strategies for patients with NMIBC.

## Author contributions

Lars Dyrskjøt had full access to all the data in the study and takes responsibility for the integrity of the data and the accuracy of the data analysis.

### Study concept and design

Strandgaard, Nordentoft, Dyrskjøt.

### Acquisition of data

Strandgaard, Nordentoft, Birkenkamp-Demtröder, Andreasen, Prip, Lindskrog, Jensen.

### Analysis and interpretation of data

Strandgaard, Nordentoft, Salminen, Christensen, Lamy, Dyrskjøt.

### Drafting of the manuscript

Strandgaard, Dyrskjøt.

### Critical revision of the manuscript for important intellectual content

All authors

### Statistical analysis

Strandgaard, Salminen, Christensen.

### Obtaining funding

Strandgaard, Dyrskjøt.

### Administrative, technical, or material support

Andreasen, Knudsen, Rasmussen, Steiniche, Jensen, Dyrskjøt.

### Supervision

Dyrskjøt.

### Other

None.

## Acknowledgements

We thank all technical personnel at the Department of Molecular Medicine, Aarhus University Hospital, for sample handling and processing.

## Data availability

The raw sequencing data are not publicly available as this compromise patient consent and ethics regulations in Denmark. Processed non-sensitive data are available upon reasonable request from the corresponding author.

## Figure legends

Supp. Fig. S1

Detailed overview of included patients and samples. Lines represent disease courses from time of first analyzed samples to end of FU centered around the first induction course of at least five instillations of BCG (orange). Colors indicate analyses performed (tumor = sequencing of tumor material, utDNA = deep targeted sequencing of utDNA, SSB = deep targeted sequencing of SSB DNA, Protein = Urinary Olink proteomics). Shapes indicate the number of different analyses performed on samples from the same clinical visit.

Supp. Fig. S2

Clinical and immunological correlation to SSBs and utDNA. a) Kaplan-Meier plot of high grade recurrence free survival (HG-RFS) from SSB with highest number of mutations for 67 patients stratified by the level of field cancerization (split by median; log-rank test). b) Kaplan-Meier plot of recurrence free survival (RFS) from SSB with highest number of mutations for 67 patients stratified by the level of field cancerization (split by median; log-rank test). c) Kaplan-Meier plot of progression free survival (PFS) from SSB with highest number of mutations for 67 patients stratified by the level of field cancerization (split by median; log-rank test). d) Kaplan-Meier plot of post-BCG HG-RFS for 85 patients stratified by the pre-BCG level of utDNA (split by median; log-rank test). e) Comparison of pre-BCG tumor CD8 T-cell adjusted exhaustion status and post-BCG field cancerization level (Wilcoxon rank sum test).

Supp. Fig. S3

Mutation tracking across different sample types for three patients (pt 1=a+b, pt 2 = c+d, pt 3 = e+f. a,c,e: Comparison of the VAF for patient-specific mutations across different sample types and time points. X-axis = Mutated positions. Y-axis = VAF/position. Mutations analyzed were included on the panels for deep targeted sequencing. Colors indicate different sample types analyzed for each patient. b,d,f) Heatmaps showing mutation VAF of mutations in tumor WES data (T) from where mutations were selected and in normal-appearing selected site biopsies (SSBs) from multiple clinical visits (indicated by columns). Timing in relation to BCG treatment is added. BCG = Bacillus Calmette-Guérin.

