## Supplementary figures for "Field cancerization impacts tumor development, T-cell exhaustion and clinical outcomes in bladder cancer"

Supplementary figure S1

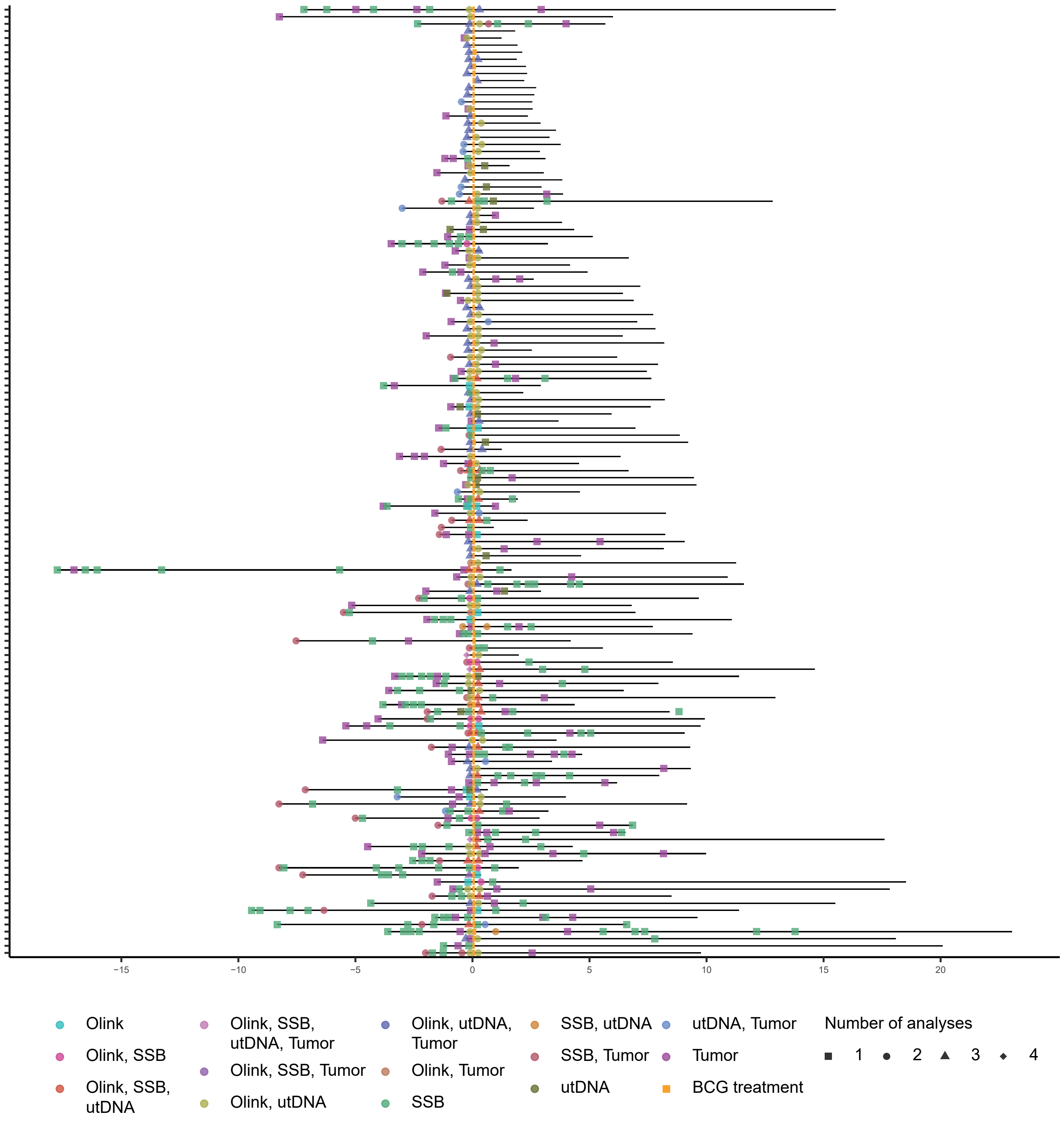


**Supp. Fig. S1**

Detailed overview of included patients and samples. Lines represent disease courses from time of first analyzed samples to end of FU centered around the first induction course of at least five instillations of BCG (orange). Colors indicate analyses performed (tumor = sequencing of tumor material, utDNA = deep targeted sequencing of utDNA, SSB = deep targeted sequencing of SSB DNA, Protein = Urinary Olink proteomics). Shapes indicate the number of different analyses performed on samples from the same clinical visit.

Supplementary figure S2

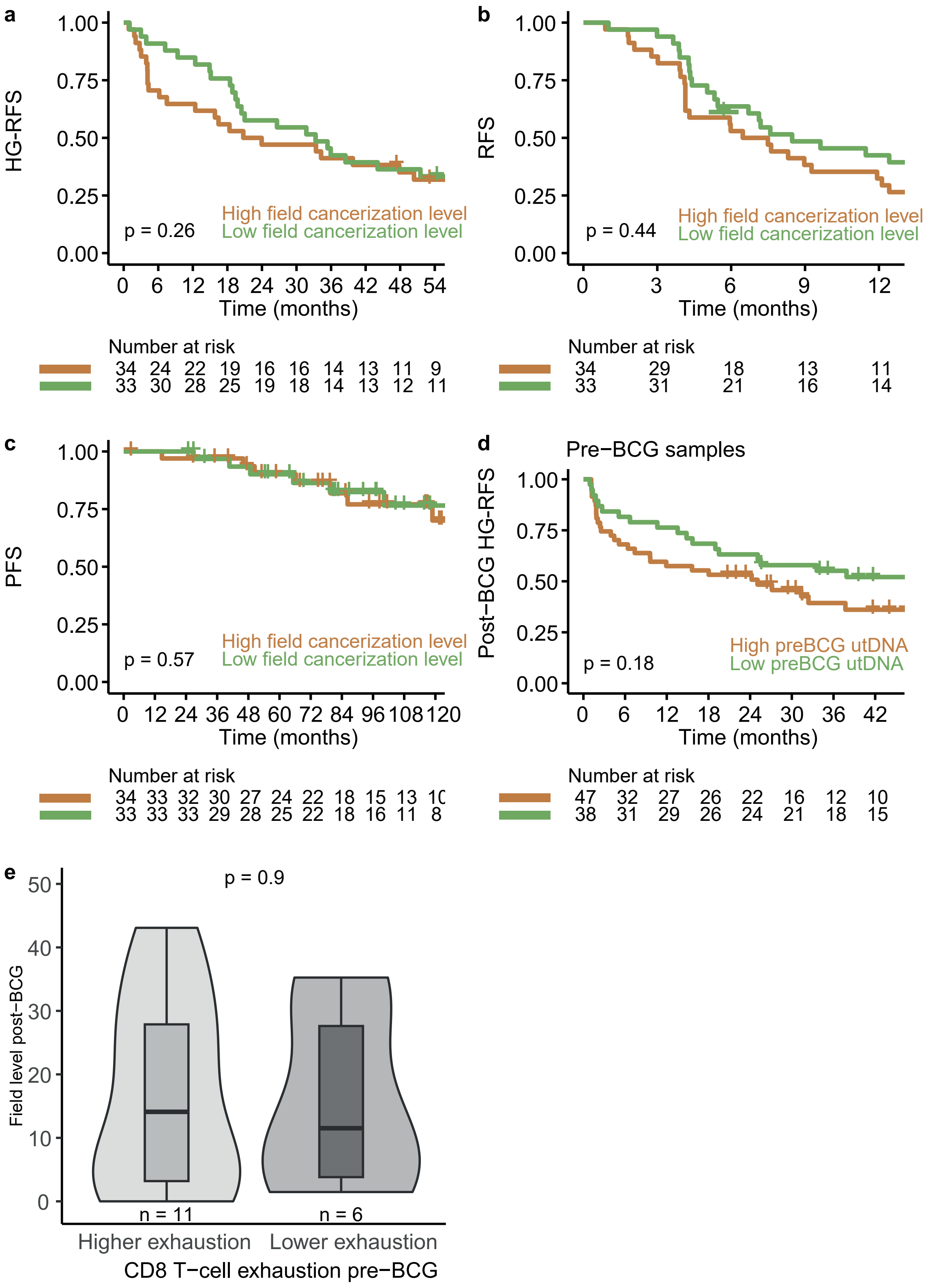


**Supp. Fig. S2**

Clinical and immunological correlation to SSBs and utDNA. a) Kaplan-Meier plot of high grade recurrence free survival (HG-RFS) from SSB with highest number of mutations for 67 patients stratified by the level of field cancerization (split by median; log-rank test). b) Kaplan-Meier plot of recurrence free survival (RFS) from SSB with highest number of mutations for 67 patients stratified by the level of field cancerization (split by median; log-rank test). c) Kaplan-Meier plot of progression free survival (PFS) from SSB with highest number of mutations for 67 patients stratified by the level of field cancerization (split by median; log-rank test). d) Kaplan-Meier plot of post-BCG HGRFS for 85 patients stratified by the pre-BCG level of utDNA (split by median; log-rank test). e) Comparison of pre-BCG tumor CD8 T-cell adjusted exhaustion status and post-BCG field cancerization level (Wilcoxon rank sum test).

Supplementary figure S3

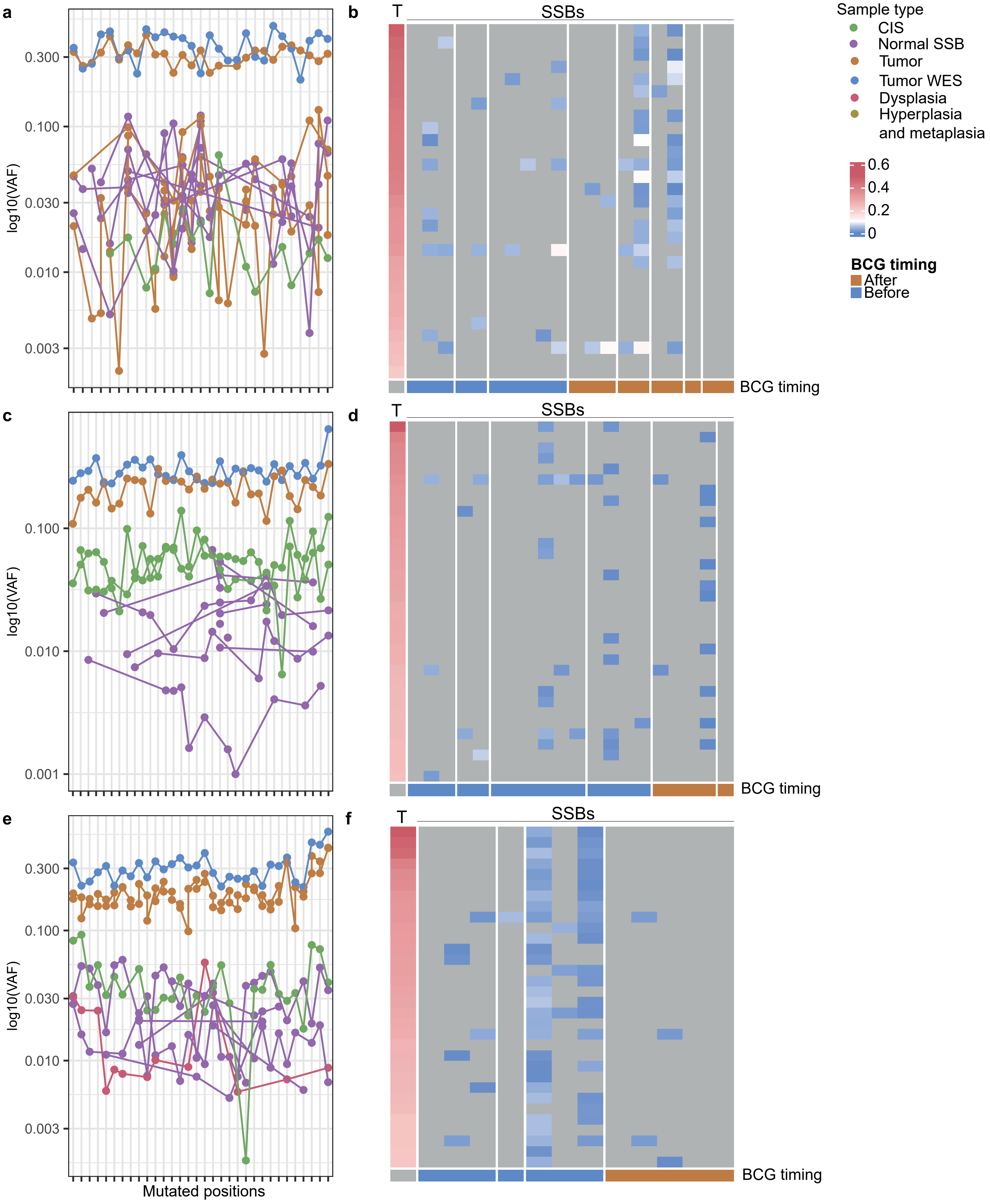

**Supp. Fig. S3**

Mutation tracking across different sample types for three patients (pt 1=a+b, pt 2 = c+d, pt 3 = e +f. a,c,e: Comparison of the VAF for patient-specific mutations across different sample types and time points. X-axis = Mutated positions. Y-axis = VAF/position. Mutations analyzed were included on the panels for deep targeted sequencing. Colors indicate different sample types analyzed for each patient. b,d,f) Heatmaps showing mutation VAF of mutations in tumor WES data (T) from where mutations were selected and in normal-appearing selected site biopsies (SSBs) from multiple clinical visits (indicated by columns). Timing in relation to BCG treatment is added. BCG = Bacillus Calmette-Guérin.
